# T-DNA integration is rapid and influenced by the chromatin state of the host genome

**DOI:** 10.1101/104372

**Authors:** Shay Shilo, Pooja Tripathi, Cathy Melamed-Bessudo, Oren Tzfadia, Theodore R. Muth, Avraham A. Levy

## Abstract

*Agrobacterium tumefaciens* mediated T-DNA integration is a common tool for plant genome manipulation. However, there is controversy regarding whether T-DNA integration is biased towards genes or randomly distributed throughout the genome. In order to address this question, we performed high-throughput mapping of T-DNA-genome junctions obtained in the absence of selection at several time points after infection. T-DNA-genome junctions were detected as early as 6 hours post-infection. T-DNA distribution was apparently uniform throughout the chromosomes, yet local biases toward AT-rich motifs and T-DNA border sequence micro-homology were detected. Analysis of the epigenetic landscape of integration showed that selected events reported on previously were associated with extremely low methylation and nucleosome occupancy. Conversely, non-selected events from this study showed chromatin marks, such as high nucleosome occupancy and high H3K27me3 that correspond to 3D-interacting heterochromatin islands embedded within euchromatin. Such structures might play a role in capturing and silencing invading T-DNA.

## Introduction

*Agrobacterium tumefaciens* is the causative agent of crown gall disease ^1–3^. Disarmed strains of *A. tumefaciens* are widely used to create genetically modified plants. *A. tumefaciens* transfers a single stranded T-DNA molecule into the plant host cell together with other virulence proteins ^1–3^. The single stranded T-DNA forms a complex with a single VirD2 protein covalently bound to its 5′. end and with several VirE2 proteins bound along the single-strand DNA. This complex is transported to the nucleus where the T-DNA integration process takes place. The T-DNA-genome junctions at the 5′ end are much more precise than at the 3′ end ^4,5^, likely owing to the role of VirD2 in protecting the 5′ end ^6^. By contrast, the frequent occurrence of DNA structural variations at the genome-T-DNA 3′ end junctions were recently shown to be due to the error-prone activity of the plant polymerase theta, a protein essential for T-DNA integration ^7^. Several lines of evidence showing that the T-DNA integrates at induced DNA double stranded breaks (DSBs) together with the typical non-homologous end-joining (NHEJ) footprints supports a model of T-DNA integration via a DSB repair pathway ^8,9^. However, integration via a double or single stranded T-DNA intermediate remains possible ^10–12^.

Several questions remain with regards to T-DNA integration: for example, the timing of integration following infection and the preferences (genetic and epigenetic) for T-DNA integration are not fully understood ^2,13,14^. The distribution of T-DNA integrations in the *Arabidopsis* genome has been examined previously, with the reports arriving at conflicting conclusions. First, a study examining over 80,000 independent integration events showed a bias for T-DNA integrations in gene-rich areas ^13^. However, these results used selective conditions and may not have been able to detect T-DNA insertions into transcriptionally inactive regions. A recent study based on the analysis of events obtained under non-selective conditions concluded that location of T-DNA integration events is essentially random ^14^. However, the relatively low number of T-DNA integration events analyzed limits the ability to identify biases in integration that may exist. These findings raise the need for an unbiased, high-throughput, system that identifies T-DNA junctions and closes the gap with the recent epigenetic data ^15–18^.

In an effort to gain an unbiased perspective of T-DNA integration, we modified the adapter-ligation mediated PCR method ^19^ from a selection based method to a selection-free method similarly to what was done recently for mapping of HIV integrations in human ^20,21^. DNA was extracted from *Agrobacterium*-infected roots at several post-infection time points. The T- DNA to genomic DNA junctions were amplified, sequenced and mapped to the *Arabidopsis* genome without the need to grow a transformed plant. Our data indicate that T-DNAs can form junctions with the genome relatively quickly (within 6 hours). Furthermore, our results show that in the absence of selection, T-DNAs integrate throughout the genome, with enrichment in regions of high nucleosome occupancy, while selected integrations occurred preferentially into hypomethylated regions with low nucleosome occupancy. In general, T-DNA insertions have some bias at the sequence level, with preferential insertions in regions that share microhomology to the T-DNA borders and with some enrichment in A-rich regions. In summary, our analysis shows that unselected T-DNA integration sites are distributed more uniformly, but not randomly, across the genome with preferences for AT-rich sequence motifs and for H3K27me3-enriched heterochromatin regions embedded in euchromatin and interacting with each other^18,22–24^.

## Results

### High-throughput detection of unselected T-DNA integrations

In order to systematically detect T-DNA integrations into the genome without the use of selection, roots of young *Arabidopsis* seedlings were cut, infected with *A. tumefaciens,* and their DNA was extracted (See Fig. 1 and details in Methods). We combined the previously described adapter ligation-PCR mediated method ^13,19^ for T-DNA junction detection with high-throughput sequencing similarly to what was done recently for HIV in human^20,21^. Due to the expected presence of deletions during T-DNA-genome junction formation via NHEJ, we used three primers located −11, −30 and −70 bp from the left border terminus.

**Fig 1.**
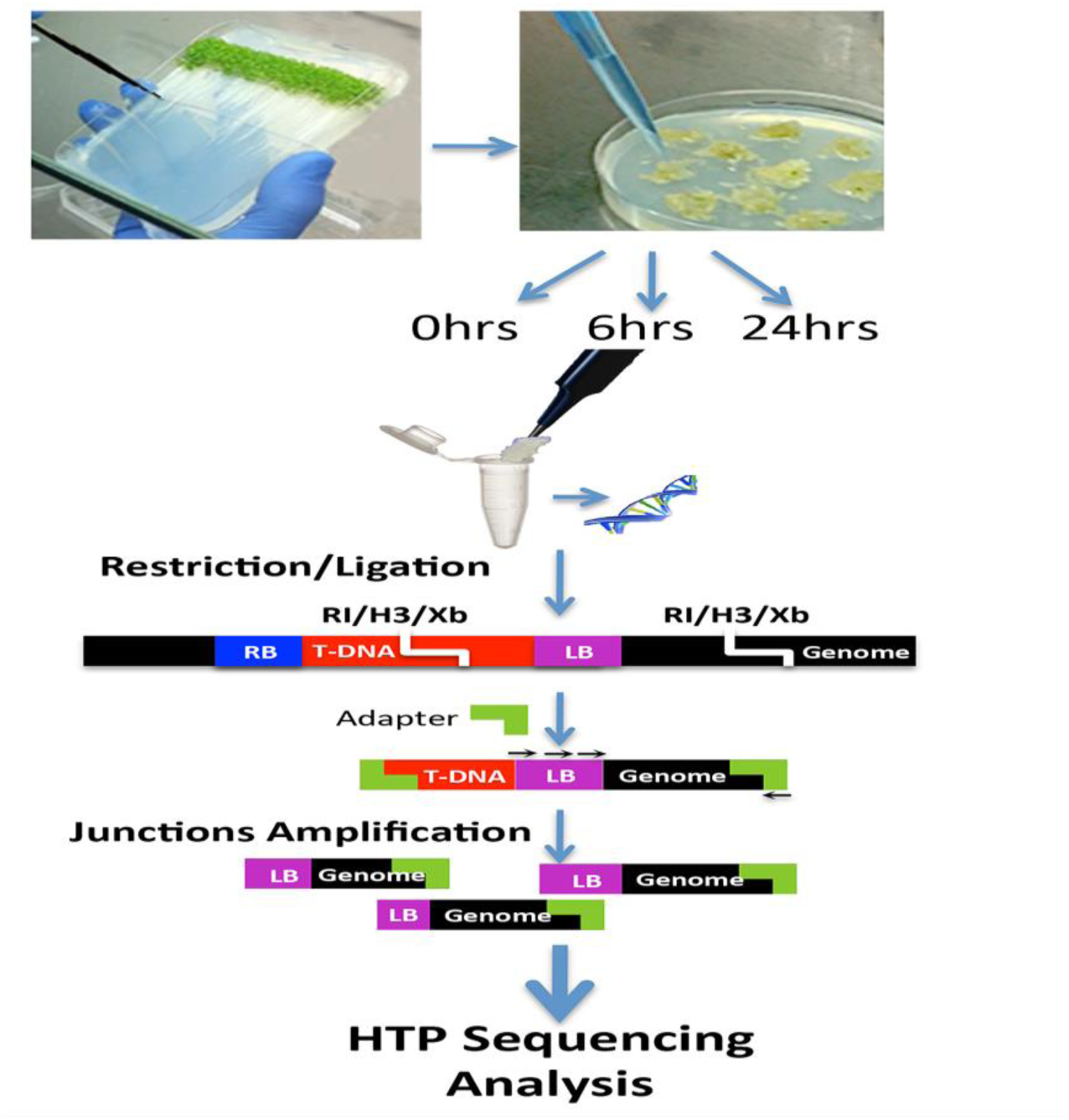
Experimental design – *Arabidopsis* roots were infected with *A. tumefaciens*. DNA was extracted at 0, 6 and 24 hours post infection. Extracted DNA was digested with 3 restriction enzymes: *Eco*RI (RI), *Hind*III (H3) and *Xba*I (Xb). An adapter was ligated to the overhang end of the digestion. T-DNA-Genomic junctions were amplified using 3 different primers from within the T-DNA and one primer from the adapter (black arrows). Amplicons were sequenced using high throughput sequencing. Adapter to adapter products were avoided as detailed in O’Malley et al. 2007 ^19^.

The high-throughput sequencing data was filtered to ensure that only high quality reads with minimal possible artifacts were used in the analysis. Only reads with a Phred score above 25 for every base were selected. Such PCR amplicon sequencing results in multiple reads of unique T-DNA junctions. We evaluated the sequences and as expected 99% of them were represented by multiple reads. The replicates were collapsed to single events after alignment. In order to avoid false positive results we used a control DNA that was extracted immediately after infection at “Time zero” (T0) with no time for integration to occur. DNA from T0 underwent the entire protocol of T-DNA junction detection, with all other experiment samples. As expected, the number of reads obtained from the T0 control prior to any filtering was only 2-3.5% the number of reads from DNA extracted 6 and 24 (T6 and T24) hours post infection (Supplemental Fig. 1). All reads from T6 and T24 that matched reads from T0 were removed from the analysis. Furthermore, T0 reads were aligned to the reference genome. Some genomic locations showed preferences for artifacts, namely if the number of reads mapped to a given location was above the mean plus two times the standard deviation, these sites were masked in the downstream analysis. Another cause for false positive results is false priming as a result of partial match between the primer and the genome. In order to remove artifacts that may result from amplification from the genome in the absence of integration, we required that every read to map to both the genome and to the T-DNA sequence. Finally, our system was based on amplification with 3 different primers from within the T-DNA. In order to avoid counting the same read more than once and to discard all the PCR duplicates, reads were collapsed so that reads from the same primer with junctions mapping 10bp apart were counted only once. In total, we identified 2801 junctions likely corresponding to integration events; 1899 of these in T6 and 902 in T24.

### Distribution of selected versus unselected T-DNA integrations along chromosomes

From the 2801 unselected integrations events that we identified, the distribution of these events in chromosome four is shown in Fig. 2 (blue line and circles). There was no obvious bias of integration when considering the main chromosomal domains (centromeric, pericentric, distal, subtelomeric or telomeric). Similar results were found for the rest of the genome (Supplementary Fig. 2). By contrast, the analysis of kanamycin-selected T-DNA integrations (Fig.2 yellow/orange line) suggested that T-DNAs tend to integrate into gene rich, transposon poor, regions, i.e. with low frequency of insertion in pericentric and centromeric regions ^13^. In detail, integrations into pericentromeric regions, corresponding to 8.3% of the genome, were rare in the selection-based data, namely 2% in reanalysis of these data. In the unselected data the ratio of integration in pericentromeric regions was more than four times higher (8.2% of the events) and close to the percent of pericentromeric regions in the genome, consistent with the expected value for random integration.

**Fig 2.**
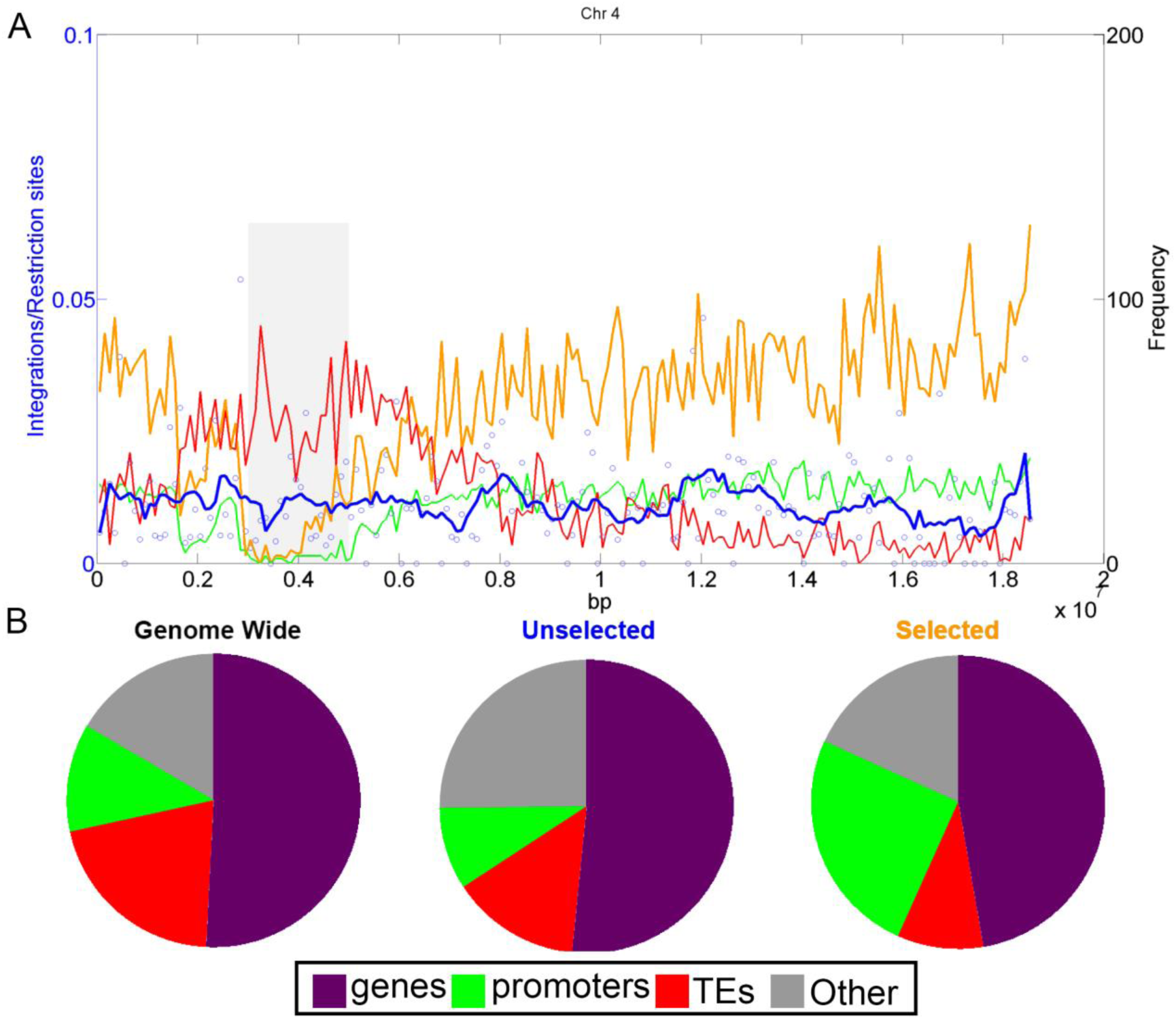
Association of genomic features with selected and unselected T-DNA integrations. A –The genomic distribution of unselected and selected T-DNA integrations across Chr 4. The numbers of T-DNA integrations (circles, and smoothed blue line) do not show correlation with the distribution of TE (red line) and promoters (green line). T-DNA integration under selective conditions (orange line) correlates with genes/promoters ^13^. B- The portion of each genomic feature: TE (red), genes (purple), promoters (green) and the remaining regions (Other (grey)). The portion of genomic features is represented across all the genome (genome wide) and according to the amount of T-DNA integration: without selection (Unselected) and with selection (Selected- data from Alonso et al., 2003 ^13^). Selected events ^13^ show enrichment in promoters (χ^2^ test, compared to unselected events, p = 2.88E-24) and a decrease in TE regions (χ2 test, compared to unselected events, p = 0.005).

Since the original publication of Alonso et al (2003) ^13^ there has been much improvement in the genome annotation as well as with epigenetic data. Therefore, we reanalyzed the |80,000 selected integration events with updated genome annotation, TAIR10, in a single base resolution, characterizing the genomic features at the site of the integration. Alonso *et al*. (2003) proposed that integrations occur in genic regions ^13^ which we confirmed (Fig. 2A). In addition, we showed that selected integrations data is biased toward promoters (green line) (Fig. 2B). By contrast, the unselected data did not show biased integration into genes, promoters or transposable elements (TE, red line) (Fig. 2A and B)

### Sequence bias at sites of unselected T-DNA integration

We did not find any bias for or against the GC content in single, di, or tri nucleotides. This confirms and extends the data from Kim et al. 2007, who reported only minor bias of GC content at the integration sites of unselected events ^14^. To further investigate the possible effect of sequence on the T-DNA integration we used two algorithms for sequence motif analysis, HOMER ^25^ and MEME ^26,27^. The genome was divided into non-overlapping bins of 400bp. Bins that contain at least one integration event were used for the analysis while random sampling of the genome was used as control. Overall, 2328 sequences of 400bp length each were used as a dataset for sequence investigation.

Some sequence motifs were found to be significantly enriched at the integration regions (Fig. 3). The most significant motif found to be enriched by both tools (P-value = 1e-147, HOMER, E-value=1.7e-234, MEME) was a motif whose consensus, CACCAC, matches to the left border of the T-DNA (Fig. 3). This motif is present in 25 percent of the input sequences (585 bins). The motif can be indicative of microhomology-mediated integrations that are frequently observed during non-homologous end-joining events ^28^ and during T-DNA integration ^4,5,29^. Other motifs were found to be significantly enriched in the integration sites (P-value < 1e-21, HOMER, E-value=2.3e-483, MEME), and these motifs tend to be AT-rich (Fig. 3).

**Fig 3.**
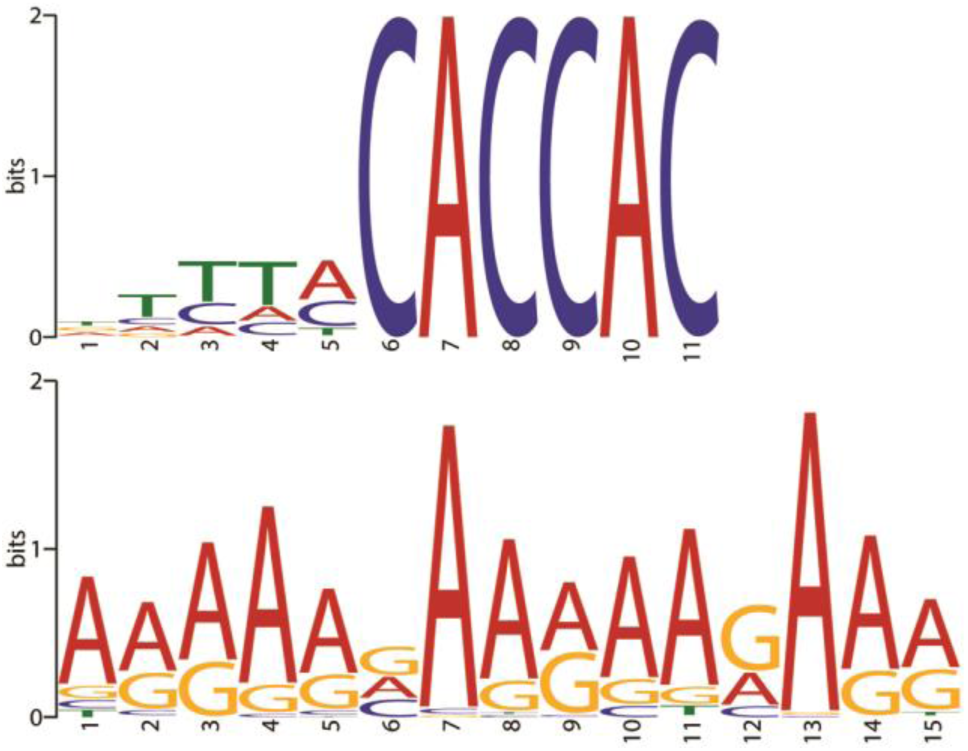
Sequence motifs associated with integration sites. The motifs CACCAC (P-value = 1e-147, HOMER, E-value=1.7e-234, MEME) and A-rich (P-value < 1e-21, HOMER, E-value=2.3e-483, MEME) were associated with integration sites.

As positive controls all the three consensus motifs of the restriction enzymes GAATTC of EcoRI (P-value = 1e-119), TCTAGA of XbaI (P-value = 1e-112), and AAGCTT of HindIII (P-value = 1e-46), integral part of the adapter ligation method (Methods), were found by HOMER in the motif analysis of the bins (data not shown).

### Contrasting epigenetic biases between selected and unselected integrations

Epigenetic marks are known to be involved in DNA recombination events ^30–33^. We performed a detailed investigation of epigenetic marks around T-DNA integration sites with or without selection. Since both centromeric and pericentric regions have distinct epigenetic characterization from the rest of the genome, we split the analysis for centromeric/pericentric and distal regions. We looked for the epigenetic landscape 500bp up and downstream to the integration site.

We found significant differences between selected and unselected integration events. In distal regions, DNA methylation of unselected integrations showed patterns close to random that were significantly different (p=9.21E-170, u-test) from the selected integrations which show almost zero methylation at the site of integration (Fig. 4A). In pericentromeric regions, methylation patterns were close to random and differences between selected and unselected integrations were not significant (p=0.4, u-test) (Fig. 4A).

**Fig 4.**
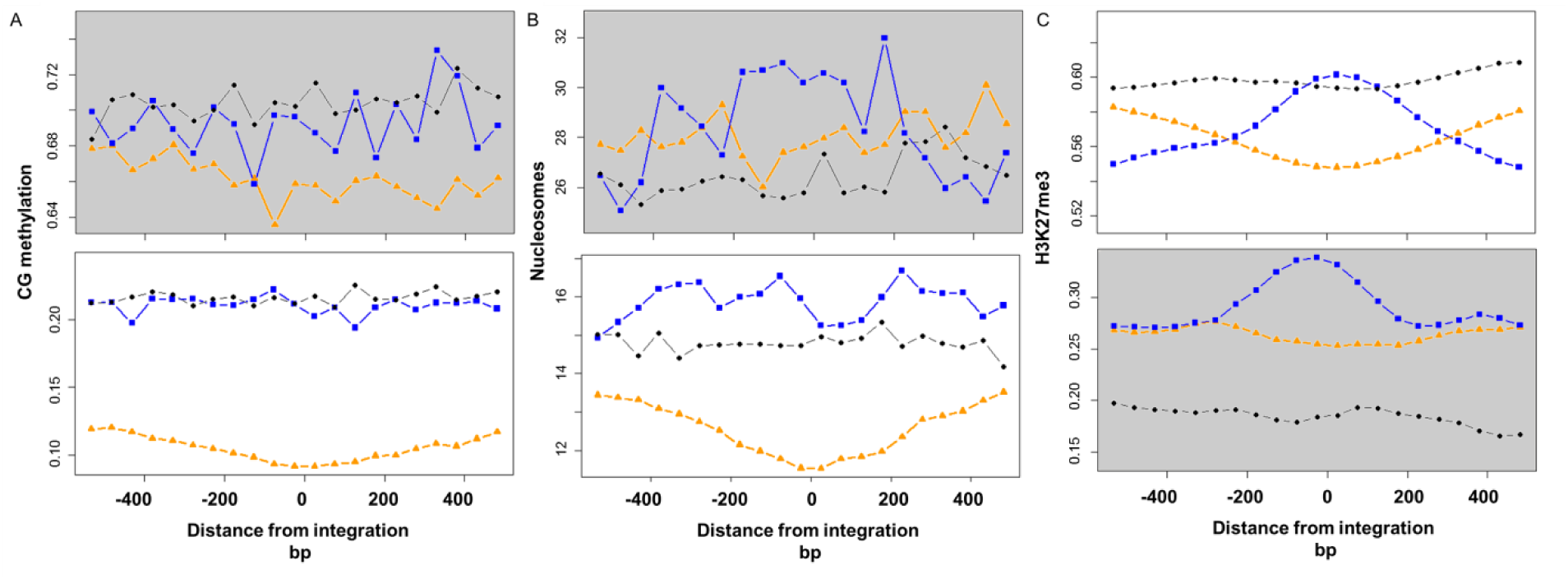
Epigenetic modifications around integration sites. The analysis was performed separately in pericentric (grey background) and in remaining (distal) chromosomal regions. The up and downstream regions to the integration point (represented as the 0 bp) are shown in the X axis. The Y axis represents arbitrary level of the epigenetic markers. Blue squares – unselected integrations. Orange triangle – integrations under selective conditions ^13^. Black circles – control, random genomic positions.–. A- CG methylation. B – Nucleosome occupancy. C- H3K27me3 modification.

Selected events showed very low nucleosome occupancy, especially at integration sites (Fig 4B). By contrast, in distal regions, unselected events showed higher nucleosome occupancy than selected events (p=1.35E-188) and slightly higher occupancy than a random dataset (p=9.97E-27) (Fig 4B). A similar trend was also observed in pericentric regions (p=1E-04, compared to selected events; p=1.12E-06, compared to control) (Fig. 4B).

In distal regions, we found a peak of H3K27me3 around the integration sites of unselected events (p<0.0035, compared to random control), while there is a “valley” for selected events (Fig. 4C). In pericentric regions, H3K27me3 levels around T-DNA insertions were overall lower than in distal regions, with higher level in selected compared to unselected regions (Fig. 4C).

## Discussion

We performed an analysis of the genetic and epigenetic landscape of T-DNA integration sites in the *Arabidopsis* genome. We reanalyzed selected T-DNA integrations ^13^ using recent data on the epigenome ^16,30^ and we performed a high-throughput analysis of unselected integrations events through the sequencing of T-DNA-genome junctions from Agro-infected *Arabidopsis* roots. We refer to these junctions as integration sites because (i) they were amplified from high molecular weight DNA, (ii) they contained hallmarks of T-DNA integration, namely, microhomologies between the T-DNA and the integration site ^34^ and (iii) only a small number of events was found at Time 0, compared to later infection stages (Supplementary Fig. 1).

### Timing

The approach used here enabled the PCR detection of T-DNA junctions independently of selection or expression of markers. Junctions were found as early as 6 hours after root infection. Earlier studies that were done without selection detected mRNA expression of a promoter-less GUS that was indicative for integrations as early as 18 h post infection in BY2 cells ^35^. Since mRNA transcription initiation, elongation, accumulation to a detectable amount of transcript and translation can delay the detection of the integration when using a reporter gene, our results provide new and direct evidence for T-DNA integration dynamics. Integration of the T-DNA in Arabidopsis roots might even occur before 6 hrs. Further work is needed to determine the exact timing of the first integrations. Another open question is the number of integrations across the time line. In our data we see reduction in the amount of integration between T6 to T24. This counterintuitive observation may result from various biological reasons, such as a stress response in the cut roots or increased apoptosis of infected root cells over time.

### Sequence integration biases

Regarding integration preferences, unlike previous studies that were based on selective systems, we found the global integration landscape to be unbiased toward genes, promoters or other tested genomic features. We even detected a small but significant bias toward intergenic regions (Figure 2B). At the sequence level, we found microhomology to the T-DNA left border to be involved in at least 25% of the integrations. This is probably an underestimate as we detected the CACCAC motif from the element left border but we did not consider very short (1-2 bp) nucleotides identity in the analysis. We also found that regions of unselected integrations are enriched with AT-rich motifs similar to the motif associated with meiotic recombination ^30^. It may be that this sequence bias reflects the occurrence of DNA DSBs that serve as entry points for T-DNA integration and as crossover inducers during meiosis.

### Epigenetic integration biases

The epigenetic landscape of unselected integrations differed significantly from selected integrations. T-DNA integrations under selected conditions tend to be found in “open chromatin” regions with very low cytosine methylation, low nucleosome occupancy and low H3K27me3. By contrast, without selection the T-DNA has a bias towards integration into regions with marks of heterochromatin such as high nucleosome occupancy and H3K27me3 (in particular in pericentric regions) but not of high cytosine methylation (Figure 4). Interestingly, Hi-C studies of chromatin packing have shown that such epigenetic marks (high H3K27me3 and high histone occupancy) define a chromatin state of small heterochromatin regions embedded in euchromatin that are “sticky”, namely, highly interacting regions ^18,22–24^. We speculate that such structures might serve as “landing” sites for incoming T-DNA—a mechanism that may protect the genome through capturing and silencing of incoming DNA into regions prone to breaks or nicks ^24^. It is also possible that the T-DNA-VirE2 or VirD2 complex interacts with host chromatin factors ^36,37^ that drive it to heterochromatin regions.

Our results extend an earlier study that showed the difference between selected and unselected events^14^. However, this earlier study, based on a small number of events (n = 117), reported on randomness of integration while we show a clear bias for specific genetic and epigenetic markers. What is the cause for the different integration patterns between selected and unselected events? The association of selection based integration with open chromatin is most likely due to the need of the selection marker to be expressed in order for the transformed plant to survive. It is thus reasonable to assume that the biases seen with selected events do not reflect an integration bias but rather the silencing of the transformation marker. While alternative hypotheses could be raised, there is a strong biological basis supporting silencing, as an explanation to the observed bias between selected and unselected events ^3,38–40^. This work, which provides a genome-wide analysis of genetic and epigenetic patterns of selected and unselected T-DNA integration, contributes to a better understanding of the process of T-DNA integration. It opens new prospects to study how the interaction between the incoming DNA and the chromatin structures determine patterns of integration.

## Methods

### Plants

Wild-type *A. thaliana,* Columbia-0 ecotype, seeds were surface sterilized in a solution of 30% bleach and 0.1% Triton X-100 for 10 minutes in a 50 ml conical vial (inverting every 2-3 minutes). The seeds were rinsed at least 3x in ~20 ml of sterile water. A P1000 pipette was used to transfer ~100-150 seeds onto square plates containing Gamborg’s B5 media (1.8% agarose; 20 g/L sucrose). Seeds were dispersed in a line ~3 cm from one edge of the plate. Plates were sealed with Parafilm and the seeds vernalized by placing in the dark at 4**°**C for 48-72 hours. Plates were removed from 4**°**C and placed in a growth chamber at 22**°**C with constant light in an upright position so that the roots grow down along the surface of the media. The seedlings were allowed to grow for 10-12 days before infecting.

### Bacteria

A frozen stock of *Agrobacteria tumefaciens* strain At1529 (GUS with intron-containing T-DNA binary vector pBISN1 in strain A348; Narasimhulu *et al.* 1996 ^41^) was freshly streaked onto a YEB plate containing 50 µg/ml kanamycin and 10 µg/ml rifampicin and grown at 28**°**C for 48 hours. A 5 ml culture of YEB media (25 µg/ml kanamycin, 10 µg/ml rifampicin) was inoculated from a single *A. tumefaciens* colony and grown overnight with shaking at 28**°**C. The next morning the overnight culture was diluted 1:20 into fresh media and grown as above until an OD_600_ of ~0.8 was reached. The bacteria were pelleted at 9,000xg for 5 min, the supernatant removed and the cells resuspended in 0.9% NaCl solution. This pelleting and rinse was repeated 1 more time and the cells were diluted down to 1:100 in 0.9% NaCl.

### *Arabidopsis* root infection

Under sterile conditions the roots of the ~11 day-old seedlings were cut into 2-3 mm segments with a scalpel. Using sterile tweezers root segments were collected into bundles of 50-100 and placed on an MS plate (with 10 g/L sucrose). Root tips were not included in the bundles. Once the bundles were prepared they were inoculated with the freshly rinsed *A. tumefaciens* bacteria, enough bacterial culture was added to cover the root bundles entirely. After 15 minutes the excess bacteria and liquid were removed gently with a pipette. Root bundles for the 0-time point were transferred to a microcentrifuge tube, rinsed 3x in 0.9% NaCl and quickly frozen in liquid nitrogen. The plates containing the remaining root bundles were sealed in Parafilm and incubated at 22**°**C in the dark. At 6 hours and 24 hours post-infection root bundles were removed from the plate, transferred to a microcentrifuge tube and washed 3x in 0.9% NaCl, frozen in liquid nitrogen and stored at −80**°**C. At 48 hours post-infection one or two root bundles were stained for GUS expression as described in Zhu *et al*., 2003 ^42^. In most cases 70-90% of cut root ends in a bundle were positive for GUS staining. If the percentage of GUS-positive root segments was lower than 70% the experiment was terminated because of the low transformation efficiency.

### Arabidopsis DNA extraction

Root tissue from each time point was processed in parallel and care was taken to keep samples separate from one another in order to reduce cross contamination. The lyophilized root bundles were physically disrupted using a Qiagen TissueLyzer set at 30 Hz for 2 min with 30 nm tungsten-carbide beads. Total genomic DNA was isolated from the root tissue using the Sigma Gene-Elute kit (G2N70-1KT) according to the manufacturer’s protocol. The 200 µl eluted genomic DNA was ethanol precipitated using 100 µl of 7.5 M ammonium acetate and 600 µl of ice cold >95% ethanol and spinning for 20 min at 0**°**C. The DNA pellet was washed once with 70% ethanol, air-dried for ~10 min, resuspended in 10 µl of TE buffer and stored at 4**°**C. To isolate the high molecular weight DNA from lower weight DNA that may contain unintegrated T-DNA, the total DNA was loaded onto a 0.7% agarose gel and run at 90 volts for 1 hour. The band containing the high weight DNA was excised with a scalpel and the DNA extracted from the gel fragment on glasswool treated with Sigmacote in a nested microcentrifuge tube column. The DNA-containing liquid extracted from the column was ethanol precipitated as described above and resuspended in 20 µl of TE buffer. The DNA was stored at 4**°**C for short-term storage (<48 hours) and −80**°**C for longer storage.

### Adapter ligation-mediated PCR

The adapter ligation-mediated PCR is based on the protocol described by O’Malley *et al*., 2007 ^19^. Their published protocol was developed to identify T-DNA junctions in stably transformed, clonal *Arabidopsis* plants using Sanger sequencing methods to identify and map the T-DNA-flanking regions to the *Arabidopsis* genome. We adapted their protocol to identify T-DNA-flanking regions from a population of *Arabidopsis* root cells containing potentially thousands of independent T-DNA integration events spread throughout the *Arabidopsis* genome. Additionally, instead of generating a clone library from a PCR amplicon and sequencing independent inserts, we used the Illumina HiSeq2000 sequencer platform to sequence the amplicons directly after the adapter ligation-mediated PCR. The O’Malley et al., 2007 protocol was followed with some modifications as described here. The primers used to generate the adapters and for subsequent PCR are shown in Table 1 Adapters with specific overhangs for EcoRI, HindIII or XbaI were generated by annealing the long and short adapters as follows: mix 20 µl of 5 µM long strand with 20 µl of 5 µM short strand adapters for either EcoRI, HindIII or XbaI adapters in 1,210 µl of 1 mM Tris, pH 8.3, in a microcentrifuge tube. Vortex the tubes and place it in a waterbath at 96**°**C for 2 min and allow to cool to room temperature over 30 minutes.

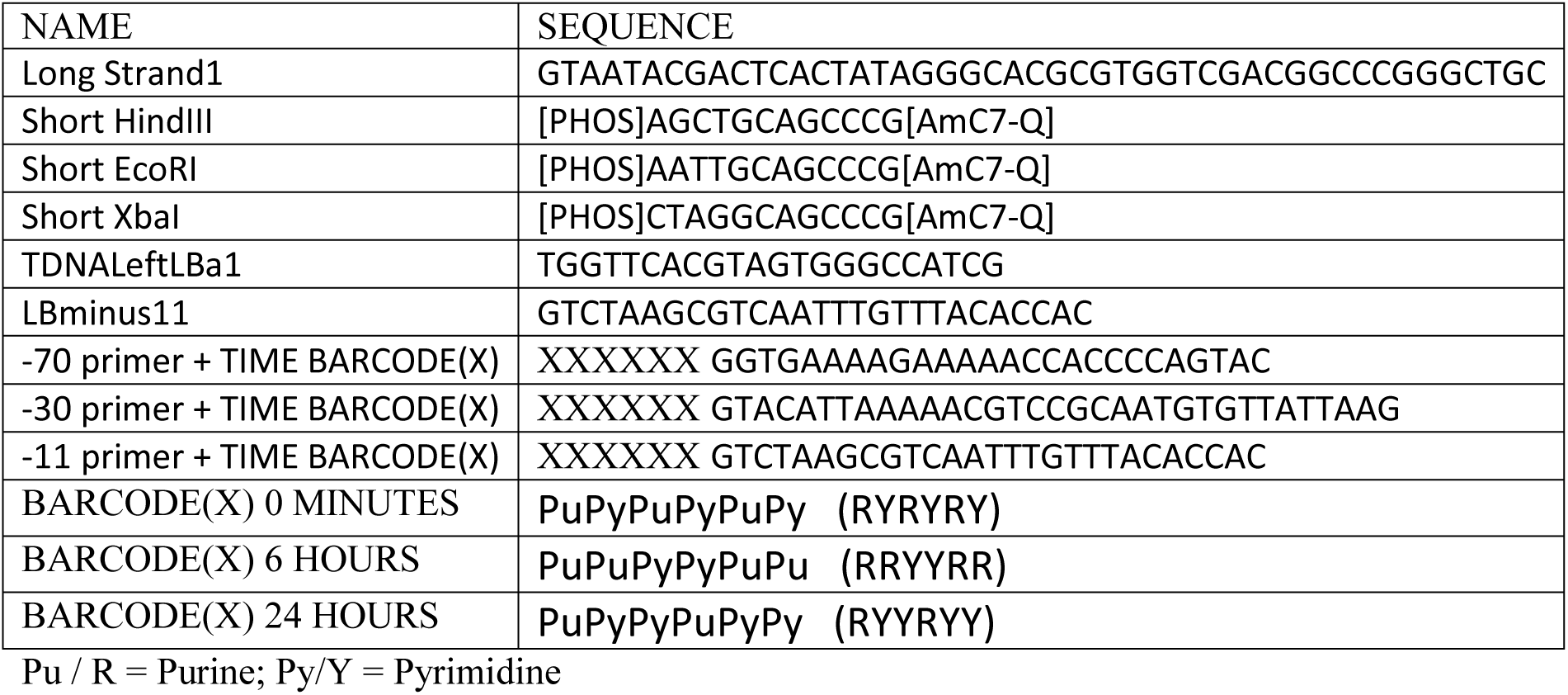
TABLE 1

For each time point, high molecular weight DNA from three or four independent root infections was pooled to create a source of template DNA for adapter ligation-mediated PCR. Each root infection time point used 4-5 bundles of roots, with each bundle containing up to 100 root segments from dozens of *Arabidopsis* seedlings, so that template DNA from each time point contained potentially thousands of independent T-DNA integration events. From the pooled high molecular weight DNA, for each time point separately, 30-50 ng of the DNA were subjected to a combined digestion and ligation process. The DNA was incubated with 2 units each of EcoRI, HindIII and, XbaI, along with 10 units of T4 DNA ligase, 0.25 µl of each adapter in NEB ligation buffer in a 40 µl volume. The combined digestion-ligation reaction was allowed to incubate at room temperature overnight. The following day the first round PCR reaction was set-up using 8 µl of the digestion-ligation product as a template in a thin-walled 200 µl PCR tube. In addition to the template the 20 µl PCR reactions contained 2 µl of 10X PCR buffer, 0.8 µl of 10 mM dNTPs, 1 µl of the LBa1 primer, 1 µl of the AP1 adapter primer, 0.1 µl Taq polymerase (either Sigma Taq of Takara) and 7.1 µl of PCR grade water. The tubes were placed in the thermal cycler and run for 10 cycles at 98**°**C for 20 seconds, 72**°**C for 2.2 minutes, then for 15 cycles at 96**°**C for 20 seconds, 67**°**C for 2.2 minutes. For the second round of PCR, 0.5 µl of the first round reaction was used as a template with the same set-up as before, but instead using 1 µl of the nested LB primer and 1 µl of the nested AP2 adapter primer. The tubes were placed in a thermal cycler and run first for five cycles at 96**°**C for 20 seconds, 88**°**C for 20 seconds, and 72**°**C for 45 seconds, and then for 23 cycles at 96**°**C for 20 seconds, 62**°**C for 20 seconds and 72**° HiSeq2000** C for 45 seconds. Because we intended to sequence the PCR amplicons and not generate clone libraries we made a number of changes to the nested LB primers. We used three LB primers with 3’ terminal base at positions, −11, −30 and −70, respectively, relative to the canonical LB T-DNA cleavage site. Each of these three versions of the LB primer were also generated with different barcodes, one specific for each time point, that would allow us to pool reactions for Illumina sequencing (see Table 1). The decision to use a set of three LB primers at increasing distance from the canonical cleavage site is because often the LB is imperfectly processed, resulting in deletions that can exceed 100 bp. However, with the Illumina short read sequencing technology we could not use a single LB primer set well back from the LB cleavage site because in many cases sequencing from the LB in the absence of any LB deletion would not allow us to sequencing far enough into the adjacent *Arabidopsis* DNA to map the location in the genome. By using three separate primers at varying distances from the LB cleavage site we increased our odds of identifying junctions that occurred with the expected LB processing and also those that might be formed in cases where up to ~65 bp of the LB had been removed in the T-DNA integration process. In the case of both the round-one and round-two PCR reactions, 5 µl were loaded onto a 1% agarose gel and run to check for amplification. In all cases, no product is visible in the first-round reactions, but in the second-round reactions a smear of DNA amplicons can be seen extending from ~50 bp to 500 bp. This smear is expected because we are amplifying thousands of independent T-DNAs junctions and each junction can be of a varying size depending on how near the adapter ligated to the left border integration position. Two or three independent round-two PCR reactions were run and the products from each time point combined and the concentration determined using a NanoDrop 1000 instrument. Equal amounts of DNA from each time point of the second round PCR products were pooled and then cleaned using a Qiagen MinElute kit. The pooled amplicons were quantified by NanoDrop and 30 µl of ~10ng/µl of DNA was used for Illumina sequencing.

### Illumina sequencing

Sequencing was performed in the INCPM center at the Weizmann Institute using the TruSeq ChIP Libary Preparation Kits and HiSeq2000 sequencer.

### Data filtering and alignment

To ensure the use of high quality reads, raw read from Illumina sequencer were filtered using PRINSEQ ^43^. Only reads with Phred score above 25 for every base were used for the downstream analysis. Reads that matched T0 reads were removed from the other time points (T6, T24) using CD-HIT ^44^. Out of the remaining reads only reads which contained one of the primers used from within the T-DNA, plus 4 bases downstream to the primer were used for the analysis. The remaining reads were mapped to TAIR10 *Arabidopsis* genome using blast ^45^. The position of the best alignment was chosen for every read. Positions that mapped +/- 10 from each other and originate from the same primer were collapsed to one. Finally, T0 reads were mapped to the genome and divided to bins of 400bp each, locations which showed preference to T0 positions, higher than mean plus two times the standard deviation, were excluded from the analysis.

### Data analysis

#### Sequence motif analysis

The genome was divided to no overlapping bins of 400bp. Bins which contained at least one integration were used for the analysis of the unselected dataset. Selected integration were taken from TAIR10 annotation files based on publicly available data ^13^. Motifs analysis was done using MEME ^27^. Another motif analysis was done using HOMER ^25^. In HOMER integration dataset was tested versus the whole genome.

#### Association with genomic features and epigenetic marks

TAIR10 genomic features were used for the genomic features analysis. Epigenetic data were provided by Assaf Zemach and were described by ^16^. The epigenetic data were binned every 50 bp. The mean values 500 bp up- and downstream of each of the integrations were calculated for all the occurrences in a certain region.

## Acknowledgements

We would like to thank Dana Averbuch and Veronika Berman for their help in the data analysis. This work was supported by an ERC grant (TRACTAR) to AAL.

